# Small molecule inhibition of multiple RNA binding proteins underlies Musashi-2 independent phenotypes

**DOI:** 10.1101/2022.09.20.508735

**Authors:** Kathryn Walters, Marcin Piotr Sajek, Aaron Issaian, Amber Baldwin, Evan Harrison, Elisabeth Murphy, Miles Daniels, Julie Haines, Kirk Hansen, Angelo D’Alessandro, Neelanjan Mukherjee

## Abstract

RNA binding proteins (RBPs) are key regulators of gene expression. Small molecules targeting these RBP-RNA interactions are a rapidly emerging class of therapeutics for treating a variety of diseases. Ro-08-2750 (Ro) is a small molecule inhibitor identified as a competitive inhibitor of Musashi(MSI)-RNA interactions. Here we show Ro potently inhibits adrenocortical steroidogenesis and viability independent of MSI2 in multiple cell lines. We identified Ro-interacting proteins using an unbiased proteome-wide approach and discovered it is broadly targeting RBPs. To confirm this finding, we leveraged the large-scale ENCODE data and found a subset of RBPs whose depletion phenocopies Ro inhibition. We conclude that Ro is a promiscuous inhibitor of multiple RBPs, many containing RRM1 domains. Moreover, we provide a general framework for validating the specificity and identifying targets of RBP inhibitors in a cellular context.

## INTRODUCTION

RNA binding proteins (RBPs) contain RNA-binding domains (RBDs) that bind to specific regulatory elements in mRNAs to control splicing, export, decay, translation, and localization^1^. These regulatory interactions are dysregulated in many human diseases including neurological disorders to cancer^2^. Despite driving disease states, RBD-RNA interfaces have not been historically targeted for small molecule inhibitors^3^. Recently, however, RBPs have emerged as a novel space for small molecule development and substantial effort has been made to the Musashi family of proteins, in particular^4,5^.

In mammals, the Musashi family of RBPs contains two homologous members: Musashi-1 (MSI1) and Musashi-2 (MSI2). MSI2 is critical for the development and maintenance of embryonic stem cells, hematopoietic stem cells, and neural stem and progenitor cells ^6^. MSI2 also plays important roles in several types of cancer including myeloid leukemia, breast cancer, and ovarian cancer among others^7–11^. Due to its importance in pathological states, substantial effort has been dedicated to developing MSI2 inhibitors^4^. Unlike most RBPs, there are several small molecules that act as competitive inhibitors of MSI2:RNA binding^12–15^. In addition, ω-9 monounsaturated fatty acids are allosteric inhibitors of MSI2:RNA binding^16^. The small molecule Ro-08-2750 (Ro) was identified as a competitive inhibitor of MSI-RNA interactions from a large chemical screen^17^. Recent structural and biochemical characterization concluded that it binds to the RRM1 domain of MSI2 to displace target mRNAs^12^. Ro treatment leads to reduction in disease burden in a murine AML leukemia model, inhibited replication of SARS-CoV-2, and improved muscle dysfunction in myotonic dystrophy type 1^12,18,19^. In all these contexts, the Ro-induced phenotypes were presumed to be dependent on MSI2, though most studies did not directly test MSI2 dependence.

MSI2 is important for fertility in steroid producing mouse gonadal tissue, as well as, steroidogenesis in human adrenocortical cells^20–22^. Here we show that Ro is a potent inhibitor of human adrenocortical steroidogenesis. However, inhibition of steroidogenesis and other Ro-dependent phenotypes were not rescued by a previously characterized Ro-resistant MSI2^12^. Using orthogonal unbiased experimental strategies, we identified a subset of RBPs, many with RRM1 domains, that both interact with and phenocopy Ro. Altogether, our findings indicate that Ro is a promiscuous inhibitor of multiple RBPs, but not MSI2.

## RESULTS

### Ro inhibits steroidogenesis in a dose-dependent manner

Angiotensin II (AngII) stimulation of immortalized adrenocortical H295R cells is a robust model for human steroidogenesis^23,24^. We previously found that siRNA depletion of MSI2 in AngII-stimulated H295R cells led to decreased aldosterone production^22^, indicating that MSI2 promotes steroidogenesis. Therefore, we hypothesized that Ro-mediated inhibition of MSI2 would also reduce aldosterone production. Indeed, Ro inhibited aldosterone production in a dose-dependent manner with an IC50 of 1.50 uM +/− 0.154 (Figure 1A). Furthermore, Ro also inhibited cortisol production with an IC50 of 0.682 uM +/− 0.056 (Figure 1B). At higher concentrations, Ro caused a decrease in cell viability and proliferation (Figure 1C-D, respectively). Ro-dependent decreases in aldosterone, cortisol, and viability were also observed in unstimulated cells (Supplemental Figure 1A-B). These observations indicated that Ro potently inhibits human steroidogenesis upstream of glucocorticoid and mineralocorticoid synthesis.

**Figure 1:**
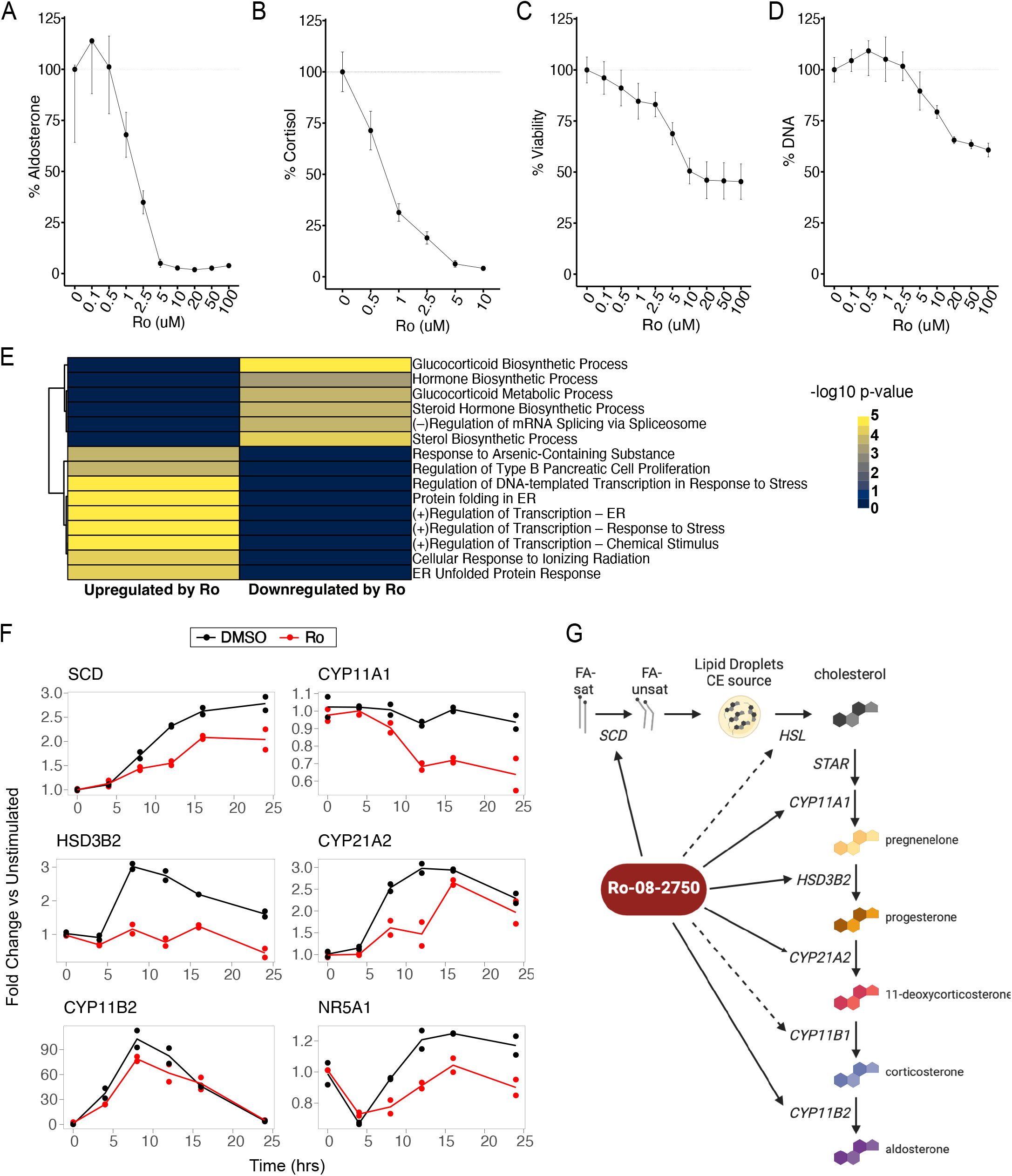
Ro inhibits steroidogenesis. Ro dose dependent effects on the percent A) aldosterone, B) cortisol, C) viability, D) cell number relative to DMSO treated H295R cells that were stimulated with 10nM Ang II. (aldosterone) or 10uM Forskolin (cortisol). E) Heatmap of the statistical significance (LRT, padj <.05) of the overlap of genes downregulated or upregulated in response to Ro with gene ontologies. F) Fold change of mRNA levels compared to unstimulated controls for DMSO (black) or Ro (red) treated H295R cells at multiple times post-AngII stimulation (10nM). G) Model for how Ro modulates steroidogenic gene expression.

### Ro coordinately downregulates multiple steps of steroidogenic gene expression

To investigate Ro-mediated inhibition of steroidogenesis, we performed an AngII stimulation RNA-seq time course (six time points in duplicate) in H295R cells treated with either Ro or vehicle (DMSO) and detected 1206 genes with significant expression differences (Supplemental Table 1). The most consistent and strongest signature was downregulation of transcripts encoding proteins involved in steroid and glucocorticoid production (Figure 1E, Supplemental Table 2). Ro caused coordinate downregulation of transcripts encoding enzymes involved in fatty acid from cholesterol metabolism to aldosterone synthesis (Figure 1F). Stearoyl-CoA Desaturase (SCD) is upstream of both cortisol and aldosterone production and is a plausible target for the observed broad inhibition of steroidogenesis. SCD converts saturated fatty acids to unsaturated fatty acids, promoting the production of lipid droplets that are important sources of cholesterol^25^. Ro treatment attenuated AngII-mediated induction of SCD mRNA and protein (Figure 1F and Supplemental Figure 1C). We also found that the mRNA encoding NR5A1, a transcription factor that controls the expression of many of the steroidogenic genes, was downregulated in the presence of Ro (Figure 1F). Taken together, we conclude that Ro coordinately represses multiple steps of the steroidogenic gene regulatory program, including fatty acid and cholesterol metabolism (Figure 1G and Supplemental Figure 1D).

### Multiple Ro phenotypes in different cell lines are MSI2 independent

We next verified that downregulation of the steroidogenic pathway upon Ro treatment was dependent on MSI2. We reasoned that expression of Ro-resistant MSI2 would rescue the loss of aldosterone production. Detailed biochemical analysis from earlier studies identified a mutation in the first RRM1 domain of MSI2 (R100A) that prevented Ro binding, but retained the ability to bind RNA targets^12^. We created a stable, doxycycline inducible FLAG-MSI2-R100A H295R cell line and found that FLAG-MSI2-R100A did not rescue Ro-mediated decreases in aldosterone or cell viability (Figure 2A, 2B, and Supplemental Figure 2B). Cell-type specific differences could have been responsible for the inability of FLAG-MSI2-R100A to rescue the Ro-mediated phenotypes in H295R cells, so we turned to K562 cells in which earlier studies demonstrated a Ro-dependent decrease in cell viability^12^ that we recapitulated (Supplemental Figure 2C). However, neither expression of FLAG-MSI2-R100A or FLAG-MSI2-WT was able to rescue Ro-mediated cell death (Figure 2C and Supplemental Figure 2C-D). The lack of rescue could not be explained by variability of dox induced FLAG-MSI2-R100A protein expression (Figure 2D). These data indicate that, unexpectedly, multiple Ro-mediated phenotypes across different cell lines are independent of MSI2.

**Figure 2:**
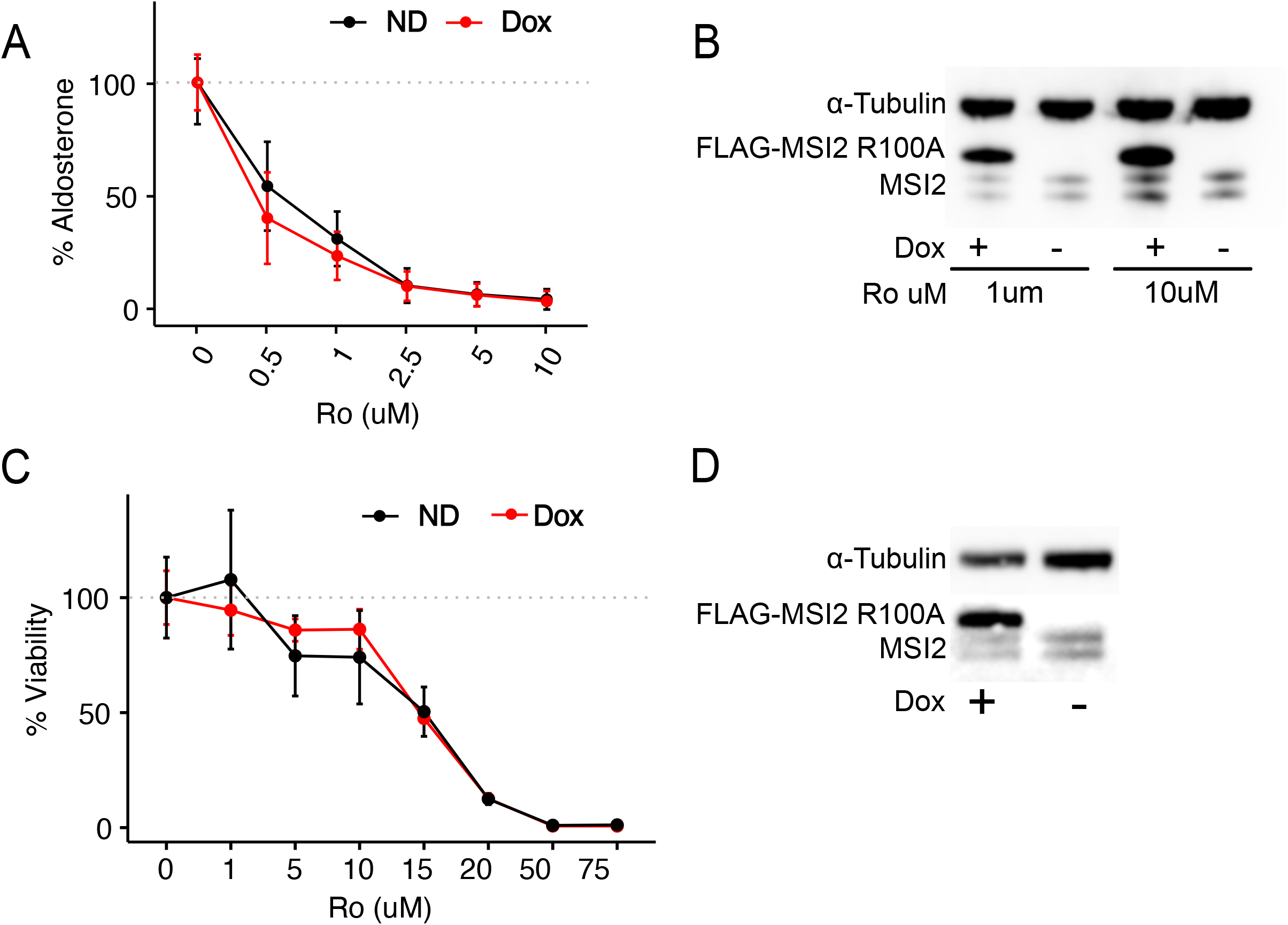
Ro phenotypes are MSI2 Independent. A) Ro dose dependent effects on the percent aldosterone relative to DMSO treated H295R-R100A cells with or without doxycycline (5 ug/mL). B) Western blot of FLAG-MSI2-R100A and endogenous MSI2 in Ro treated cells with or without doxycycline. α-Tubulin included as a loading control. C) Ro dose dependent effects on the percent viability measured relative to DMSO in K562 cells transfected with FLAG-MSI2-R100A plasmid with or without doxycycline. D) Western blot of FLAG-MSI2-R100A and endogenous MSI2 at two doses of Ro in K562 cells with or without doxycycline. α-Tubulin included as a loading control.

### Ro binds and may inhibit other RRM1 containing RBPs

Since Ro treatment caused cellular phenotypes that were not dependent on MSI2, we sought to identify Ro-interacting proteins in an unbiased manner using a Proteome Integral Solubility Alteration (PISA) assay^26^ (Figure 3A). We performed a one-dimensional PISA and identified 768 proteins that exhibited a statistically significant thermal stability shift upon addition of Ro to lysates (Figure 3B). RBPs were the most highly enriched class of Ro-interacting proteins (Figure 3C, Supplemental Figure 3A, Supplemental Table 3), consistent with previous data indicating Ro binds to RRM domains^12^. Nearly 20% of the Ro-interacting proteins were RBPs (145/768). In contrast, MSI2 exhibited a statistically significant, yet very modest thermostability shift (Figure 3B). Many of the RBPs identified contain RRM1 domains (Figure 3D), raising the question of whether the RBPs bound by Ro were also inhibited by Ro. If that were true, we reasoned that depletion of such RBPs would generate similar transcriptomic signatures to Ro treatment. For the 29 RBPs that were both PISA interactors and had shRNA knockdown profiles in the ENCODE compendium^27^ (Figure 3E, venn diagram), we calculated the correlation between Ro induced fold changes and shRNA knockdown induced fold changes for the 1181 genes that were differentially expressed upon Ro treatment. For the plurality of Ro-interacting RBPs, shRNA depletion had a statistically significant positive correlation with Ro treatment (Figure 3E). MSI2 depletion exhibited no correlation with Ro treatment, consistent with our earlier data indicating MSI2 independent function of Ro (Figure 3E). These data pointed to an alternative model where Ro binds to multiple RBPs in a dosage-dependent manner. Such inhibition of RBP-binding would lead to alterations in the balance between RNA, RBPs, and RBP-RNA complexes that could promote stress granule assembly^28–31^. We observed dose-dependent stress granule formation upon Ro treatment (Supplemental Figure 3B). Altogether, we conclude that Ro inhibits steroidogenesis and cell viability independent of MSI2, but dependent on preferential binding and inhibition of RBPs and other RRM1 containing RBPs.

**Figure 3:**
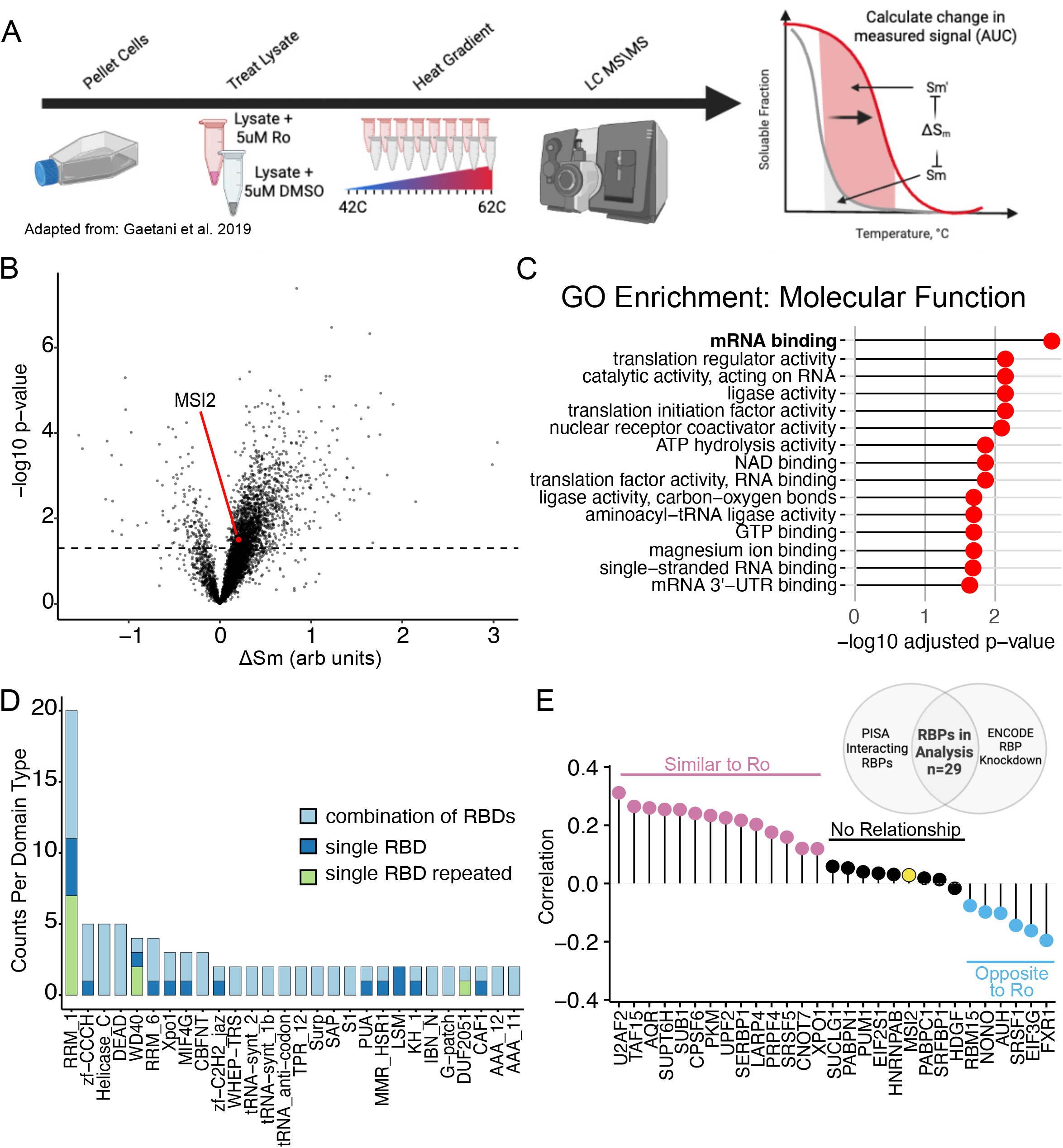
Ro interacts with multiple RBPs. A) Depiction of PISA assay (adapted from^26^). B) Volcano plot of the change in relative temperature-dependent solubility of measured proteins with respect to Ro treatment compared to -log(10) p-value. C) Top 15 gene ontology molecular functions enriched in proteins identified as Ro interactors from PISA assay. D) Bar plot showing the count of each RBD domain type for RBPs enriched as PISA interacterors. E) Correlation between gene expression changes in Ro treated H295R cells and gene expression changes from RBP knockdown in K562 cells (ENCODE) for the subset of genes with Ro-dependent expression changes (see methods). RBPs colored by the significance of the correlation coefficient (positive = pink, not significant = gray, and negative = blue).

## DISCUSSION

Ro has profound regulatory effects on the process of steroidogenesis and cell viability that are not dependent on MSI2. In contrast to previous studies, we find that these phenotypes are not rescued by expressing a modified MSI2 (R100A) that binds RNA but not Ro. The biochemical properties of the R100A and other mutations altering RNA and Ro-binding were determined via in vitro binding assays that showed MSI2-R100A could rescue Ro-mediated deficiencies in colony formation of mouse MLL-AF9 bone marrow cells^12^. An important difference between this study and our analyses is assay timing; colony formation assays take weeks, while our cell viability and steroid production assays take only 2-3 days. Because our assays are short in duration, they are more likely to register direct, rather than secondary, effects. Other studies also indicate that MSI2 inhibitors may be independent of MSI2. For example, Ro promotes cell death in HCT116 cells upon treatment with putative MSI2 inhibitors Ro or (-)gossypol with or without MSI2 knockout^13^. Rescue experiments with MSI2 drug resistant mutants or small molecule inhibitors in cells without MSI2 are reliable, *directed* approaches to establish target dependence; yet are limited by their low throughput and inability to provide alternative targets. Creation of knockout cells is similarly limited in that it is time consuming, lacks feasibility for essential genes, and does not provide detailed biochemical information. To address these limitations, we employed unbiased and orthogonal strategies to identify Ro-interacting proteins using (PISA) and potential functional RBP targets by correlating Ro-mediated and RBP-dependent gene expression (ENCODE data). Based on these data, we propose that Ro broadly targets RBPs and RRM1-containing RBPs. In addition, our findings define a general experimental framework for validating the specificity and identifying targets of other RBP inhibitors in a cellular context.

While Ro is unlikely to be a specific inhibitor of MSI2, it still elicits phenotypic changes that can be used to treat a variety of diseases. The initial study characterizing Ro as a MSI2 inhibitor showed that in an aggressive murine MLL-AF9 murine leukemia model, 19 days of Ro treatment led to reduction in disease progression^12^. Recent studies that identify MSI2 as a protein of interest from biochemical or phenotypic screens utilized Ro to support the association of the observed phenotype with MSI2. In one case, MSI2 was identified as upregulating the microRNA (*miR)-7* which prevents excessive autophagy. Autophagy is a contributor to muscle wasting which is a symptom of myotonic dystrophy^19^. When myotonic dystrophy type 1 fibroblasts were treated with Ro, *miR-7* was also upregulated, phenocopying the MSI2 regulation^19^. MSI2 was identified as an RBP interacting with the SARS-CoV-2 genome and subsequent experiments showed Ro treatment led to a modest decrease in SARS-CoV-2 nucleocapsid production, which can be used as a surrogate for viral growth^32^. Neither of these studies performed additional rescue or validation experiments testing the MSI2-dependence of the Ro phenotype. Another study also found that Ro decreased SARS-CoV-2 nucleocapsid production, however in this case the putative target is the viral papain-like protease^18^. We found that Ro inhibits steroidogenesis doses lower than those that affect cell viability (Figure 1A-D). Therefore, Ro or further refined analogues could be valuable therapeutics for treating primary aldosteronism and Cushing’s syndrome, which involve excessive production of aldosterone or cortisol, respectively. Additionally, Ro may have therapeutic potential for adrenocortical carcinomas and other cancers. Molecular classification of adrenocortical carcinomas revealed that the class with high steroidogenic capacity had the worst survival outcomes^33^. It would also be interesting to test if Ro were able to promote the repression of cholesterol biosynthesis, which is important for TP53-mediated liver tumor suppression^34^. Thus, Ro may have important therapeutic potential in a variety of diseases.

Both synthetic and natural small molecule inhibitors have been identified for RBPs (ELAVL1, LIN28, and MSI2) that regulate subsets of RNAs via specific interactions with distinct RNA regulatory elements^12,13,35–38^. Inhibitors have also been developed for core components of the spliceosome, such as SF3B1 and U2AF^39–41^. Risdiplam and Branaplam, which are used to treat spinal muscular atrophy, presumably bind to the RNA-protein interface formed by the SMN2 pre-mRNA and the U1 snRNP^42,43^. Small molecular modulation of translation targeting the ribosome and core translational machinery has led to compounds targeting EIF4A and EIF4E^44,45^. Despite these successes, the strong electrostatic affinity of the RNA binding pocket and the sequence/structural conservation of these RBDs amongst RBPs, makes it challenging to identify highly specific small molecules that are competitive inhibitors of RBPs^5^. Furthermore, many small molecule inhibitors have micromolar affinity for their RBP target and are likely to bind to other off-target proteins in cells, as we show here for Ro; implying that in a cellular context this class of small molecule inhibitors may drive pleiotropic phenotypes. Since Ro inhibited steroidogenesis at a lower concentration than cell viability in H295R cells (Figure 1A-D), it would be quite informative to determine the Ro dose-dependent differences in interaction partners using 2D PISA, which interrogates solubility shifts across multiple drug concentrations^26^. Since RBPs and RNA regulatory elements have emerged as promising targets for modulation by small molecule inhibitors, studies like ours can establish target specificity and identify new inhibitor targets.

## METHODS

### Cell Culture

H295R cells were cultured in complete media (DMEM/F12 media, 10% Cosmic Calf Serum, 1% Insulin-Transferrin-Selenium) at 37°C, 5% CO_2_. For all ELISA assays, cells were cultured in low sera media (DMEM/F12 media, 0.1% Cosmic Calf Serum, 1% Insulin-Transferrin-Selenium). K562 cells were cultured in RPMI10 media (RPMI 1640 +L-Glutamate, 10% Fetal Bovine Serum).

### Aldosterone/Cortisol ELISA, Presto Blue and FluoReporter Assay

H295R cells were plated in complete media at a density of 20,000 cells/well or 10,000 cells/well respectively for either aldosterone or cortisol experiments, respectively. Twenty-four hours after plating, complete media was removed and replaced with 110uL low sera media containing Ro-08-2750 (Tocris) suspended in DMSO or DMSO as vehicle control, and 10nM Angiotensin II (Sigma-Aldrich) for stimulation or water for unstimulated cells. For the cortisol experiments, cells were stimulated with 10uM forskolin (TCI Chemicals). After 24 hours, either 100uL or 20uL supernatant was collected for aldosterone or cortisol competitive ELISA (Invitrogen), respectively. When applicable, supernatants were diluted to a final volume of 100uL using low sera media and the ELISA was performed according to the manufacturer’s instructions. The remaining media was removed, and cells received 100uL of complete media with 1:10 dilution of PrestoBlue Cell Viability Reagent (Invitrogen). This was incubated for 2 hours at 37°C and then read on a Synergy HT plate reader (BioTek). All media was removed from the cells and the plate was frozen at −80°C. Upon thawing, the Fluoreporter Blue Fluorometric dsDNA Quantification kit (Invitrogen) assay was run according to the manufacturer’s instructions and read on a Synergy HT plate reader. Four parameter logistic (4PL) curve was fitted to a standard curve and used to calculate aldosterone/cortisol concentrations. IC50 values were calculated using the drc R package^46^.

### RNA Sequencing

H295R cells (500,000 cells per well) were plated in complete media. After 24 hours, complete media was removed and replaced with low sera media for an additional 24 hours. Cells were then stimulated with 10nM AngII and treated with DMSO control or 5 uM Ro. Cells were harvested at 0, 4, 8, 12, 16, 24 hours post-stimulation. Cells were collected in TRIzol and RNA was isolated using DirectZol Miniprep Plus kit (ZYMO) with on-column DNase I digestion. Isolated total RNA was quantified using the Qubit RNA Broad Range assay kit (Invitrogen) with the Qubit 3.0 fluorometer. Poly(A) RNA was enriched from 1ug of total RNA using the NEBNext poly(A) mRNA magnetic isolation module (New England Biolab). At the final elution step of polyA selection, the RNA-bound beads were resuspended into 1X Fragment, Primer, Elute (FPE) buffer from the KAPA RNA Hyper Prep kit (Roche), heated at 85°C for 6 minutes for fragmentation. The resulting RNA was used as input into library preparation at the first strand synthesis step using the KAPA RNA Hyper Prep Kit (Roche). Libraries were sequenced to a depth of 20 million paired-end 2×150bp reads each on the NovaSeq 6000 (Illumina) at University of Colorado Genomics and Microarray Core.

### RNA-seq Analysis

Salmon^47^ was used for quantifying transcript levels from all libraries using Gencode v26 (parameters -l A --allowDovetail --validateMappings). All downstream analysis was performed in R. Briefly, Salmon data were imported using the tximport library^48^. Differential gene expression was performed using DESeq2 library^49^ with likelihood ratio test (padj < 0.05). Pathview library^50^ was used for pathway data visualization.

### Western Blot analysis

Cells were collected in 1x Laemmli buffer containing 5% beta-mercaptoethanol. Samples were heated at 95°C for 5 minutes to denature and then loaded onto a 15 well, 4 to 12% gradient Bis-Tris, 1.0–1.5 mm Novex Mini Protein Gel (Invitrogen). Samples were transferred onto an iBlot mini NC stack nitrocellulose membrane (Invitrogen) using the iBlot2 with P0 transfer program (Invitrogen). Membranes were blocked with 5% non-fat milk in 1X tris-buffered saline with 0.1% tween (TBST) at room temperature for 30 minutes to 1 hour. Membranes were sequentially incubated with primary antibodies (all 1:2000 in 5% milk TBST) and HRP-labeled secondary antibodies (1:10k in 5% milk TBST) (Supplemental Table 4). The signals were detected by chemiluminescence using the Azure Biosystems Sapphire biomolecular imager.

### K562 Viability Assay

K562 cells (2 million cells in 100 uL) were transfected with 1ug of plasmid (MSI2-WT or MSI2-R100A) using the Neon Transfection System (Invitrogen, 1,450, 10 ms, 3 pulses). The transfection was recovered into 10mL of complete media and then split evenly into untreated or 5ug/mL doxycycline treated conditions. From each condition, 10,000 cells/well were plated in a 96-well plate. Ro +/− dox media was added to each well to give a final concentration of 5ug/mL dox and (0, 1, 5, 10, 15, 20, 50, 75 uM) Ro in 200uL complete media. The remaining cells were plated for western collection in untreated and doxycycline treated conditions. Cells were grown for 24hr at 37C in 5% CO_2_. 100uL of cells were transferred to an opaque white bottomed plate and 100uL of Cell-Titer Reagent (Promega) was added. After a 15-minute room temperature incubation, luminescence was measured on a Synergy HT plate reader (BioTek).

### Plasmid Creation

pRD-RIPE plasmid was digested with AgeI and BstXI. FLAG-MSI2-R100A and FLAG-MSI2-WT were obtained from Twist Biosciences and amplified to contain homology ends for the digested pRD-RIPE (Supplemental Table 5). Gibson homology was used to insert the MSI2 fragments into the pRD-RIPE plasmid and transformed into DH5a competent *E. coli* generated from the Mix and Go Transformation Buffer kit (ZYMO). Plasmids were sequenced for validation.

### Stable Cell Line Creation

A loxP-flanked blasticidin resistance cassette was inserted into the AAVS1 safe harbor locus of H295R cells via CRISPR/Cas9 as has been done previously^51^ (Supplemental Figure 2A). Once the H295R LoxP cell line was established, MSI2 pRD-RIPE plasmids were electroporated into the cell line using the Neon electroporation system (Invitrogen, 1100V 30ms and 2 pulses) and allowed to grow at 37°C for one week. The cells were treated with puromycin (5ug/mL) for one week, allowed to recover, then treated again for another week. Cells were validated using PCR and western blot and screened for steroidogenic induction using aldosterone ELISAs (Invitrogen).

### Proteome Integral Solubility Alteration

The PISA method was performed as previously described^26^. In brief, lysates from cells were adjusted to 1 mg/mL and divided into 120 40 μL aliquotes, 60 treatment and 60 control. Treated lysate samples were incubated with up to 100 μM ligand. Samples were heated in a thermocycler (LifeEco, Bioer) at various temperatures (42, 42.4, 43.3, 45, 47.2, 50, 53.4, 56.3, 58.5, 60.4, 61.4, 62 °C) Samples were isobaric labeled with TMT reagent (TMT10plex, Thermo Fisher Scientific) as recommended by the manufacturer. Peptide fractionation was performed on a Gemini NX-C18 column (Phenomenex) using a Dionex UltiMate 3000. Peptides were separated using the following gradient: 3% B (0–16.5 min), 3–45% B (16.5–41.5 min), 45–65% B (41.5–46.5 min), 65–100% B (46.5–48.5 min), 100% B (48.5–53.5 min) with solvent A (10 mM ammonium formate, pH 10.0) and solvent B (10 mM ammonium formate, 75% ACN, pH 10.0) at a flow rate of 0.5 mL/min. TMT labeled peptides were analyzed by nano-ultrahigh performance (UHP)LC–MS/MS (Easy-nLC1200, Orbitrap Fusion LumosTribrid, Thermo Fisher Scientific). Proteome Discoverer 2.2 (Thermo Fisher Scientific) was used for the database search and TMT quantification. Quantification results were log2 normalized and the ΔS_m_ value was determined for every protein. Statistical analysis was performed by Perseus^52^.

### GO and KEGG enrichment on PISA data

We used clusterProfiler R library^53^ to perform GO and KEGG enrichment on PISA interacting proteins. We used a false discovery rate for multiple testing correction with a significance cutoff of 0.05. The list of human RBPs and domains were taken from^54^.

### Immunofluorescence

H295R cells were plated on poly-D-lysine coated coverslips. Cells were treated with DMSO, 1, 5, or 10uM of Ro for 24 hours. Cells were then fixed for 10 minutes in Neutral Buffered Formalin solution. Cells were blocked and permeabilized in CAS-T (CAS-Block (Thermo Scientific) with .2% Triton-X) for 30 minutes followed by incubation for 1 hour at room temp with anti-HuR/ELAV1 antibody (1:000 dilution in CAS-T) (Supplemental Table 4). Cells were washed with PBS-T (PBS with 0.1% Tween) and incubated for 40 minutes at room temp with Anti-mouse Alexa Fluor^®^ 488 antibody (1:1000 dilution in PBS-T) (Supplemental Table 4). Coverslips were mounted on slides using Vectashield mounting media containing DAPI and imaged at 60× magnification using a Deltavision Elite widefield fluorescence microscope (GE).

### Correlation between Ro expression changes and RBP knockdown expression changes

The ENCODE consortium generated RNA-seq data for shRNA depletion of > 200 RBPs in K562 cells. We downloaded the kallisto transcript quantification data of shRNA depletion RNA-seq for the subset of RBPs that were in the ENCODE database and were PISA interactors (n=29, Supplemental Table 7) as well as control shRNAs from K562. Data were imported using tximport R library^48^. Genes with counts sum >= 10 in all samples were kept for differential gene expression analysis. Differential gene expression was performed using the DESeq2 library^49^ with Wald test. We applied the same analysis to the H295R RNA-Seq from 12hr Ro-treated cells. Namely, transcripts were quantified using kallisto v. 0.44^55^ (kallisto quant -i /path/to/kallisto.idx -o /path/to/outdir --plaintext read1.fastq read2.fastq). The kallisto index for gencode GRCh38 v.29 was downloaded from ENCODE^56^. Data were imported using tximport R library^48^. Genes with counts sum >= 20 in all samples were kept for differential gene expression analysis. Differential gene expression was performed using the DESeq2 library^49^ with Wald test. For the set of 1181 genes differentially expressed after Ro treatment in H295R cells and passed both expression filters, we calculated Pearson correlation coefficients between ENCODE data and Ro-treated cells using stat values.

## Supporting information

Supplemental Table 1 RNAseqDEall

Supplemental Table 2 HeatmapGOMcat all

Supplemental Table 3 GOanalysis MFall

Supplemental Table 4 Antibody Usage

Supplemental Table 5 Cloning Sequences Reagents

## Data and code availability

RNA-Seq data are available from Gene Expression Omnibus (accession number GSE213218). Processed data and code: https://github.com/mukherjeelab/2022_RoInhibitor

## Acknowledgements

We thank David Bentley, Jay Hesselberth, and Sara Johnson for the critical review of the manuscript. This material is based upon work supported by the National Science Foundation Graduate Research Fellowship Program under Grant No. 1000317291-NSFGRFP-Walters (K.W.)., the Research Experiences for Graduate and Medical Students (REGMS) Summer Fellowship Program (K.W.), Polish National Agency for Academic Exchange Bekker Program PPN/BEK/2019/1/00173 (M.P.S.), University of Colorado Anschutz Medical Campus RNA Bioscience Initiative (N.M.), Boettcher Foundation Webb-Waring Early Career Investigator Award AWD-103075 (N.M.), and National Institutes of Health 1R35GM147025-01 (N.M.). Biorender was used in Figure 1G, Figure 3A, and Supplemental Figure 2A.

## Author contributions

N.M. conceived the project; K.W., A.B., E.H., A.I., M.D., E.M., and J.H. performed experiments and collected data; K.W, M.P.S., A.I., J.H., and N.M. performed formal analysis and conducted the visualization; K.W. wrote the original draft; K.W., M.P.S., A.B., A.A., K.H., and N.M. reviewed and edited the paper; K.W., M.P.S, A.A., K.H., and N.M. acquired funding; A.A., K.H., and N.M. provided resources; N.M. supervised the project.

---

**Supplemental Figure 1:**
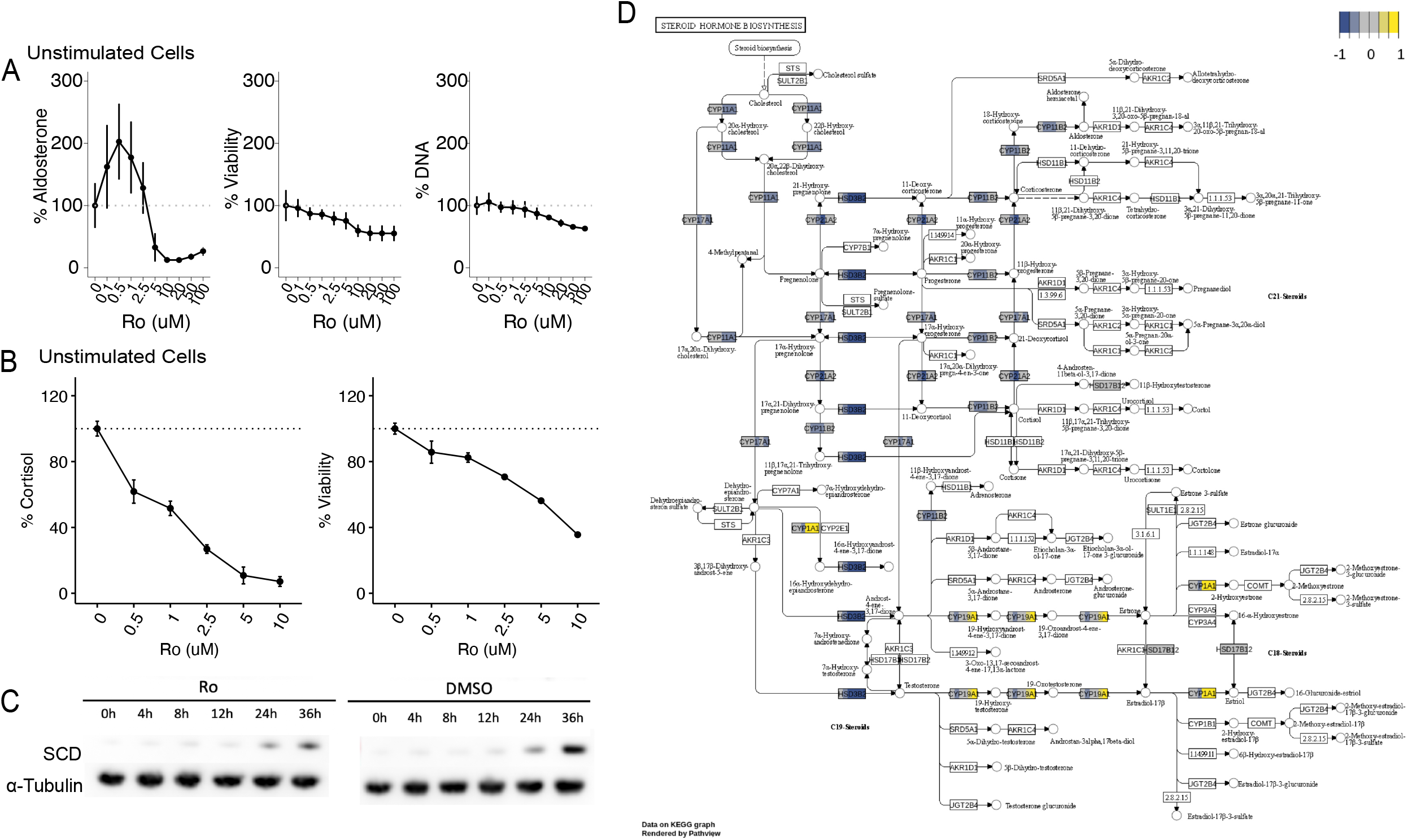
Ro inhibits steroidogenesis in unstimulated cells. A/B) Ro dose dependent effects on percent aldosterone, cortisol, viability, and cell number relative to DMSO in unstimulated H295R cells. C) Western blot of SCD protein amount in DMSO or 5 uM Ro treated H295R cells in a time course of hours post 10nM Ang II stimulation. α-Tubulin included as a loading control. D) Pathway data visualization from RNA-seq samples taken in a time course of hours post 10 nM AngII stimulation. Yellow indicates enrichment, navy indicates depletion.

**Supplemental Figure 2:**
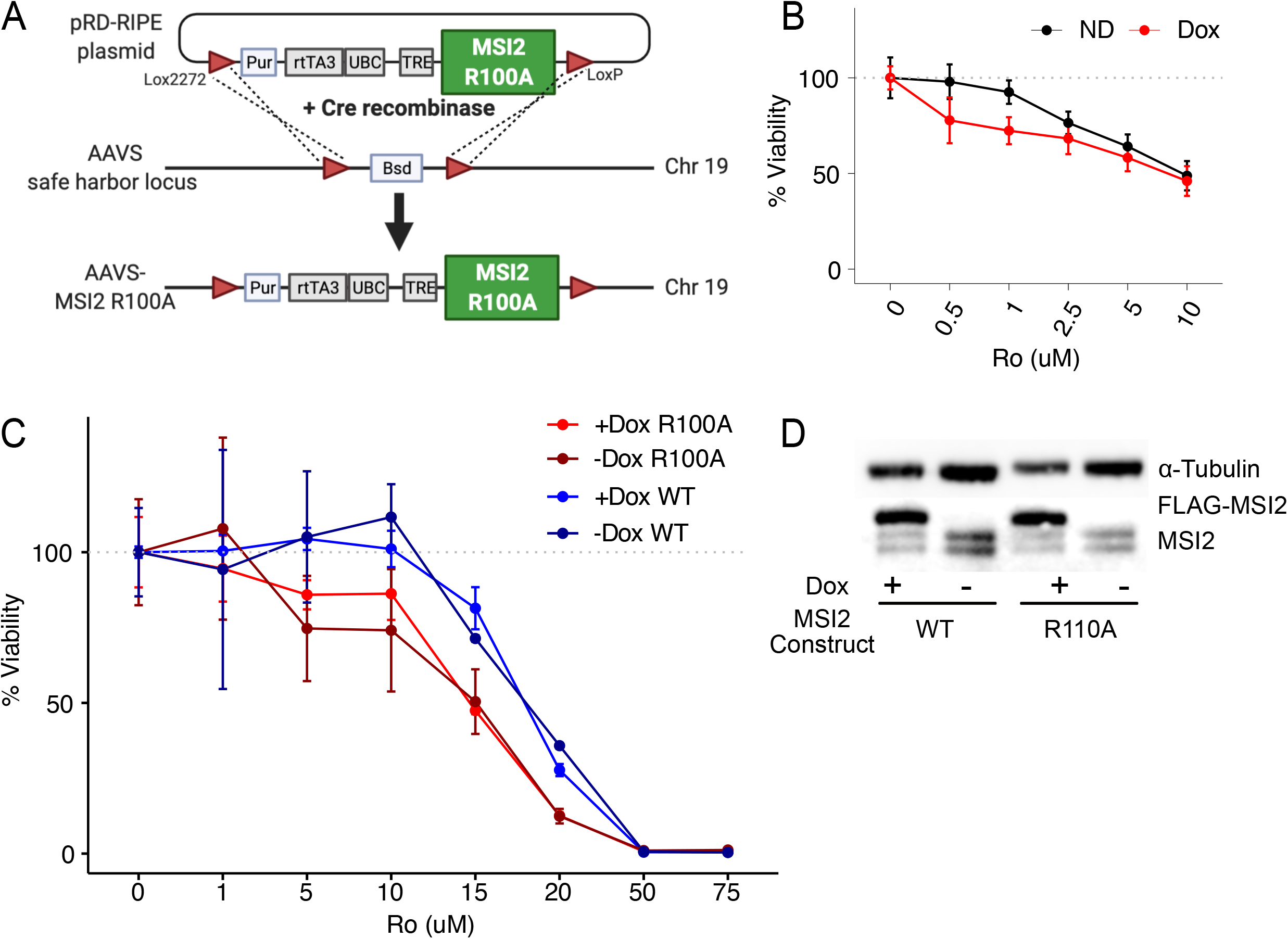
R100A MSI2 is unable to rescue Ro phenotypes. A) Diagram of the MSI2 floxed portion of the pRD-RIPE plasmid recombination into the AAVS site of H295R cells. B) Percent viability of Ro treated H295R-R100A cells, with or without doxycycline. Experiment was paired with the aldosterone data found in Figure 2A. C) Ro dose dependent effects on the percent viability measured relative to DMSO in K562 cells transfected with FLAG-MSI2-R100A plasmid or FLAG-MSI2-WT plasmid and treated with or without doxycycline. D) Western blot of FLAG-MSI2-R100A, FLAG-MSI2-WT, and endogenous MSI2 in Ro treated K562 cells, with or without doxycycline. α-Tubulin included as a loading control.

**Supplemental Figure 3:**
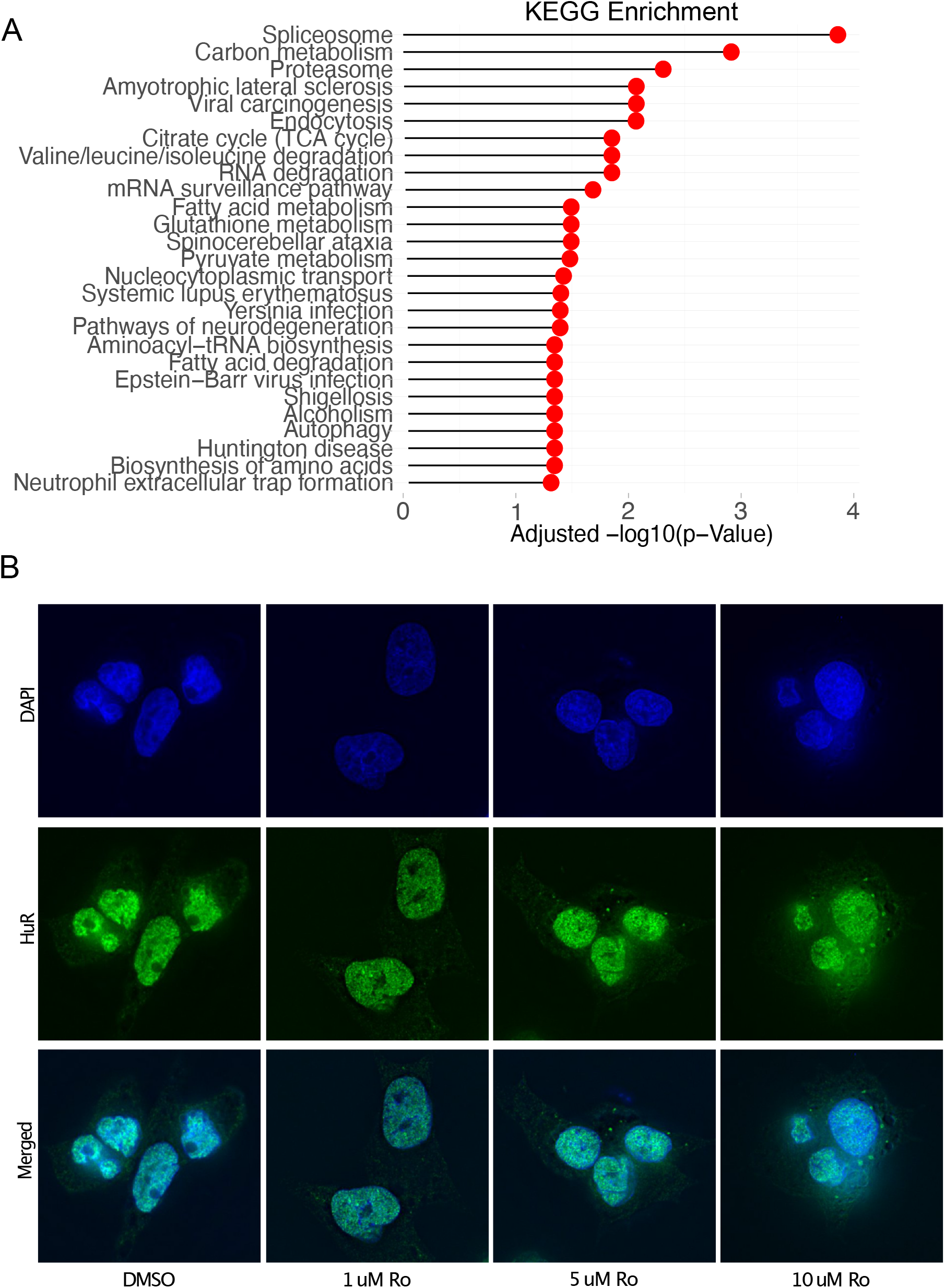
Ro treatment causes stress granule formation. A) List of significant gene ontology KEGG pathways enriched in proteins identified as Ro interactors from PISA assay. B) Immunofluorescence of a stress granule marker, HuR/ELAV1 (green), for DMSO or Ro treated unstimulated H295R cells. Nuclei are shown in blue.

## List of Supplemental Information (Found in Supplemental Folder)

Supplemental Table 1: Ro-Dependent Gene Changes

Supplemental Table 2: Full names of the Heatmap in Figure 1

Supplemental Table 3: Full names of the GO analysis in Figure 3

Supplemental Table 4: Antibody information.

Supplemental Table 5: MSI2 cloning information.

## Notes

### Competing Interest Statement

The authors have declared no competing interest.

## References

1. Keene, J. D. RNA regulons: coordination of post-transcriptional events. Nat. Rev. Genet. 8, 533–543 (2007).

2. Gebauer, F., Schwarzl, T., Valcárcel, J. & Hentze, M. W. RNA-binding proteins in human genetic disease. Nat. Rev. Genet. 22, 185–198 (2021).

3. Julio, A. R. & Backus, K. M. New approaches to target RNA binding proteins. Curr. Opin. Chem. Biol. 62, 13–23 (2021).

4. Kudinov, A. E., Karanicolas, J., Golemis, E. A. & Boumber, Y. Musashi RNA-Binding Proteins as Cancer Drivers and Novel Therapeutic Targets. Clin. Cancer Res. 23, 2143–2153 (2017).

5. Wu, P. Inhibition of RNA-binding proteins with small molecules. Nature Reviews Chemistry 4, 441–458 (2020).

6. Fox, R. G., Park, F. D., Koechlein, C. S., Kritzik, M. & Reya, T. Musashi signaling in stem cells and cancer. Annu. Rev. Cell Dev. Biol. 31, 249–267 (2015).

7. Kharas, M. G. et al. Musashi-2 regulates normal hematopoiesis and promotes aggressive myeloid leukemia. Nat. Med. 16, 903–908 (2010).

8. Ito, T. et al. Regulation of myeloid leukaemia by the cell-fate determinant Musashi. Nature 466, 765–768 (2010).

9. Kang, M.-H. et al. Musashi RNA-binding protein 2 regulates estrogen receptor 1 function in breast cancer. Oncogene 36, 1745–1752 (2017).

10. Lee, J. et al. Musashi-2 is a novel regulator of paclitaxel sensitivity in ovarian cancer cells. Int. J. Oncol. 49, 1945–1952 (2016).

11. Li, N. et al. The Msi Family of RNA-Binding Proteins Function Redundantly as Intestinal Oncoproteins. Cell Rep. 13, 2440–2455 (2015).

12. Minuesa, G. et al. Small-molecule targeting of MUSASHI RNA-binding activity in acute myeloid leukemia. Nat. Commun. 10, 2691 (2019).

13. Zhang, X. et al. Small Molecule Palmatine Targeting Musashi-2 in Colorectal Cancer. Front. Pharmacol. 12, 793449 (2021).

14. Wang, M. et al. Suppression of Musashi 2 by the small compound largazole exerts inhibitory effects on malignant cells. Int. J. Oncol. 56, 1274–1283 (2020).

15. Lan, L. et al. Natural product derivative Gossypolone inhibits Musashi family of RNA-binding proteins. BMC Cancer 18, 809 (2018).

16. Clingman, C. C. et al. Allosteric inhibition of a stem cell RNA-binding protein by an intermediary metabolite. Elife 3, (2014).

17. Minuesa, G. et al. A 1536-well fluorescence polarization assay to screen for modulators of the MUSASHI family of RNA-binding proteins. Comb. Chem. High Throughput Screen. 17, 596–609 (2014).

18. Lim, C. T. et al. Identifying SARS-CoV-2 antiviral compounds by screening for small molecule inhibitors of Nsp3 papain-like protease. Biochem. J 478, 2517–2531 (2021).

19. Sabater-Arcis, M. et al. Musashi-2 contributes to myotonic dystrophy muscle dysfunction by promoting excessive autophagy through miR-7 biogenesis repression. Mol. Ther. Nucleic Acids 25, 652–667 (2021).

20. Sutherland, J. M. et al. Developmental expression of Musashi-1 and Musashi-2 RNA-binding proteins during spermatogenesis: analysis of the deleterious effects of dysregulated expression. Biol. Reprod. 90, 92 (2014).

21. Sutherland, J. M. et al. Knockout of RNA Binding Protein MSI2 Impairs Follicle Development in the Mouse Ovary: Characterization of MSI1 and MSI2 during Folliculogenesis. Biomolecules 5, 1228–1244 (2015).

22. Fu, R. et al. RNA-binding proteins regulate aldosterone homeostasis in human steroidogenic cells. RNA (2021) doi:10.1261/rna.078727.121.

23. Parmar, J., Key, R. E. & Rainey, W. E. Development of an adrenocorticotropin-responsive human adrenocortical carcinoma cell line. J. Clin. Endocrinol. Metab. 93, 4542–4546 (2008).

24. Wellman, K. et al. Transcriptomic Response Dynamics of Human Primary and Immortalized Adrenocortical Cells to Steroidogenic Stimuli. Cells 10, 2376 (2021).

25. Paton, C. M. & Ntambi, J. M. Biochemical and physiological function of stearoyl-CoA desaturase. Am. J. Physiol. Endocrinol. Metab. 297, E28–37 (2009).

26. Gaetani, M. et al. Proteome Integral Solubility Alteration: A High-Throughput Proteomics Assay for Target Deconvolution. J. Proteome Res. 18, 4027–4037 (2019).

27. Van Nostrand, E. L. et al. A large-scale binding and functional map of human RNA-binding proteins. Nature 583, 711–719 (2020).

28. Van Treeck, B. et al. RNA self-assembly contributes to stress granule formation and defining the stress granule transcriptome. Proc. Natl. Acad. Sci. U. S. A. 115, 2734–2739 (2018).

29. Molliex, A. et al. Phase separation by low complexity domains promotes stress granule assembly and drives pathological fibrillization. Cell 163, 123–133 (2015).

30. Riggs, C. L., Kedersha, N., Ivanov, P. & Anderson, P. Mammalian stress granules and P bodies at a glance. J. Cell Sci. 133, (2020).

31. Wang, J. et al. A Molecular Grammar Governing the Driving Forces for Phase Separation of Prion-like RNA Binding Proteins. Cell 174, 688–699.e16 (2018).

32. Kamel, W. et al. Global analysis of protein-RNA interactions in SARS-CoV-2-infected cells reveals key regulators of infection. Mol. Cell 81, 2851–2867.e7 (2021).

33. Zheng, S. et al. Comprehensive Pan-Genomic Characterization of Adrenocortical Carcinoma. Cancer Cell 29, 723–736 (2016).

34. Moon, S.-H. et al. p53 Represses the Mevalonate Pathway to Mediate Tumor Suppression. Cell 176, 564–580.e19 (2019).

35. Meisner, N.-C. et al. Identification and mechanistic characterization of low-molecular-weight inhibitors for HuR. Nat. Chem. Biol. 3, 508–515 (2007).

36. Wu, X. et al. Identification and validation of novel small molecule disruptors of HuR-mRNA interaction. ACS Chem. Biol. 10, 1476–1484 (2015).

37. Wang, L. et al. Small-Molecule Inhibitors Disrupt let-7 Oligouridylation and Release the Selective Blockade of let-7 Processing by LIN28. Cell Rep. 23, 3091–3101 (2018).

38. Lan, L. et al. Natural product (-)-gossypol inhibits colon cancer cell growth by targeting RNA-binding protein Musashi-1. Mol. Oncol. 9, 1406–1420 (2015).

39. Kotake, Y. et al. Splicing factor SF3b as a target of the antitumor natural product pladienolide. Nat. Chem. Biol. 3, 570–575 (2007).

40. Kaida, D. et al. Spliceostatin A targets SF3b and inhibits both splicing and nuclear retention of pre-mRNA. Nat. Chem. Biol. 3, 576–583 (2007).

41. Chatrikhi, R. et al. A synthetic small molecule stalls pre-mRNA splicing by promoting an early-stage U2AF2-RNA complex. Cell Chem Biol 28, 1145–1157.e6 (2021).

42. Palacino, J. et al. SMN2 splice modulators enhance U1-pre-mRNA association and rescue SMA mice. Nat. Chem. Biol. 11, 511–517 (2015).

43. Sivaramakrishnan, M. et al. Binding to SMN2 pre-mRNA-protein complex elicits specificity for small molecule splicing modifiers. Nat. Commun. 8, 1476 (2017).

44. Bordeleau, M.-E. et al. Functional characterization of IRESes by an inhibitor of the RNA helicase eIF4A. Nat. Chem. Biol. 2, 213–220 (2006).

45. Moerke, N. J. et al. Small-molecule inhibition of the interaction between the translation initiation factors eIF4E and eIF4G. Cell 128, 257–267 (2007).

46. Ritz, C., Baty, F., Streibig, J. C. & Gerhard, D. Dose-Response Analysis Using R. PLoS One 10, e0146021 (2015).

47. Patro, R., Duggal, G., Love, M. I., Irizarry, R. A. & Kingsford, C. Salmon provides fast and bias-aware quantification of transcript expression. Nat. Methods 14, 417–419 (2017).

48. Soneson, C., Love, M. I. & Robinson, M. D. Differential analyses for RNA-seq: transcript-level estimates improve gene-level inferences. F1000Res. 4, 1521 (2015).

49. Love, M. I., Huber, W. & Anders, S. Moderated estimation of fold change and dispersion for RNA-seq data with DESeq2. Genome Biol. 15, 550 (2014).

50. Luo, W. & Brouwer, C. Pathview: an R/Bioconductor package for pathway-based data integration and visualization. Bioinformatics 29, 1830–1831 (2013).

51. Goering, R., Arora, A. & Matthew Taliaferro, J. RNA localization mechanisms transcend cell morphology. bioRxiv 2022.04.14.488401 (2022) doi:10.1101/2022.04.14.488401.

52. Tyanova, Temu, Sinitcyn, Carlson & Hein. The Perseus computational platform for comprehensive analysis of (prote) omics data. Nature.

53. Yu, G., Wang, L.-G., Han, Y. & He, Q.-Y. clusterProfiler: an R package for comparing biological themes among gene clusters. OMICS 16, 284–287 (2012).

54. Gerstberger, S., Hafner, M. & Tuschl, T. A census of human RNA-binding proteins. Nat. Rev. Genet. 15, 829–845 (2014).

55. Bray, N. L., Pimentel, H., Melsted, P. & Pachter, L. Near-optimal probabilistic RNA-seq quantification. Nat. Biotechnol. 34, 525–527 (2016).

56. Encff904uyq – encode. https://www.encodeproject.org/files/ENCFF904UYQ.

